# Reusing wasted bioresources for crop protection: Calcinated oyster shell powder enhances rhizospheric microbial-mediated suppression of root-knot nematodes

**DOI:** 10.1101/2025.01.14.633056

**Authors:** Qipeng Jiang, Yong Wang, Jiamin Yu, Jinfeng Wang, Lianqiang Jiang, Shiping Guo, Yu Qian, Xiangwen Yu, Dongyang Liu, Daojiang Xi, Quan Deng, Wei Ding, Shili Li

## Abstract

The root-knot nematode (RKN) *Meloidogyne incognita* is one of the most destructive plant-parasitic nematodes (PPNs) affecting crop production worldwide. Our earlier study revealed that calcinated oyster shell powder (OSP) effectively suppressed tobacco RKN disease. However, the suppression mechanism of OSP against RKNs remain unknown. Our study revealed that calcinated OSP reduced the tobacco root–knot index by more than 38% by inhibiting the migration of second-stage juveniles of *Meloidogyne incognita* (J2) in the soil. Furthermore, OSP reduced the J2 density in the tobacco rhizosphere by 43.69% and significantly increased the soil pH by 0.68; moreover, OSP increased the contents of soil exchangeable calcium (ExchCa) and exchangeable magnesium (ExchMg) by more than 50%. Moreover, soil properties, including ExchMg, ExchCa and pH, enhanced the microbial-mediated suppression of J2. Specifically, Proteobacteria and Gemmatimonadota dominated the microbial community suppressing RKN, and fungal richness contributed to the suppression of RKNs; in addition, Chloroflexi and Acidobacteria dominated the microbial community promoting RKN prosperity. Our study revealed that the reuse of wasted oyster shell powder as an innovative antagonist is a promising avenue for ecofriendly RKN management strategies.

**IMPORTANCE:** There is increased demand for ecofriendly RKN management strategies, and this study is the first to propose that the reuse of wasted oyster shell powder as an innovative antagonist against RKNs is effective, and calcinated oyster shell powder can reduce tobacco root-knot index and *Meloidogyne incognita* (J2) density by inhibiting the migration of J2s and enhancing the microbially mediated suppression of J2s in the tobacco rhizosphere. Moreover, several soil microbial and physicochemical indicators, including the contents of soil exchangeable calcium, exchangeable magnesium, pH and the relative abundance of Proteobacteria, Gemmatimonadota, Chloroflexi and Acidobacteria in plant rhizosphere were suggested to guide the development of ecofriendly PPN management strategies.

---

Root-knot nematodes (RKNs) of the genus *Meloidogyne* are among the most destructive plant-parasitic nematodes (PPNs) (1). Among the genus *Meloidogyne, Meloidogyne incognita* (*M. incognita*) causes substantial economic losses to crop production worldwide, especially in Solanaceae crop production, such as tomato (2), tobacco (3, 4) and eggplant (5). This highly obligate and soil-borne endoparasite aggregates around the plant root surface, invades plants and then establishes feeding sites, which hinders the uptake of nutrients and water, resulting in developmental retardation of plants. Moreover, RKN infection causes wounds in plant roots, which can result in further infection by other soil-borne pathogens, leading to complex plant diseases (6, 7). However, considering their safety and environmental pollution, several chemical nematicides have been banned or limited in agricultural use because of their negative impacts on environmental and human health. Hence, there is increased demand for eco-friendly products to manage RKNs.

Oyster shells are a available waste of seafood; however, they are also a good calcium-enriched natural product and alternative soil conditioner (8). In recent years, oyster shells have been widely reused as waste sources. Calcinated oyster shell powder is alkaline and contains abundant calcium ions, making it an economical product for crop production and soil amelioration. Recent studies have shown that oyster shell powder plays a positive role in increasing crop production (9), alleviating soil compaction and acidification (10, 11) and improving the fertility of soil (12). Moreover, the application of oyster shell powder to soil can potentially suppress soil-borne diseases. Shen et al. (10) reported that the addition of oyster shell powder to soil could control tobacco bacterial wilt by alleviating soil acidification and regulating the composition of the soil bacterial community. Martial (13) reported that heat-treated oyster shell powder improved the defense of Theobroma cocoa against *Phytophthora megakarya* by inducing the priming defense system and stimulating the agronomic growth of seedling plants. Nevertheless, few studies have focused on the effects of oyster shell powder on RKNs or PPNs. Notably, in a previous study, we reported that oyster shell powder effectively controlled tobacco RKN disease, but the mechanism by which oyster shell powder suppresses RKN disease remains unknown.

In general, RKN migration and invasion around plant root surfaces are affected by diverse factors, such as moisture (14), root exudates (15) and the soil microbiome (16). In recent years, culture-independent high-throughput sequencing has greatly expanded the repertoire of plant-associated microbiomes and their roles in plant disease (17, 18), allowing for us to explore the complex interactions between pathogens and other microbiomes in the disease process, and reveal the microbial indicators and microbial-mediated mechanism of disease outbreaks. Although precise and comprehensive studies have revealed the significant role of rhizospheric microbes in the suppression of RKNs (19), for example, Cao et al. reported that RKN-infected tobacco presented a richer and more diverse rhizosphere soil bacterial community than healthy tobacco did (3), works on the RKN suppression mechanism of oyster shell powder are still limited.

The objective of this study was to clarify the suppression indicators and mechanism of oyster shell powder against RKNs. The ability of oyster shell powder to control *M. incognita* growth was confirmed via pot and field experiments. To explore the mechanism by which oyster shell powder suppressed *M. incognita*, we used culture-independent high-throughput sequencing to characterize the bacterial and fungal communities in the tobacco rhizosphere. Additionally, the effects of the application of oyster shell powder on soil properties were tested. Furthermore, a structural equation model (SEM) was constructed to quantify the effects of soil properties and rhizospheric microbial community on *M. incognita*.

## Results

### Effects of oyster shell powder on tobacco root–knot nematode infection in pot experiments

The effects of oyster shell powder (OSP) treatment on root-knot nematode (RKN) infection in tobacco roots were investigated in pot experiments (Fig. 1), and the results revealed that OSP markedly suppressed RKN infection and migration in soils and promoted tobacco growth and root activity (Fig. 1C). Specifically, the giant cells and *M. incognita* J2 (J2) in tobacco roots significantly decreased with increasing OSP content in the soils (Fig. 1A), and the migration of J2 in the soils mixed with 0.4% OSP was significantly lower than that in the CK (Fig. 1B, Fig. S1). The root-knot indices of tobacco roots significantly decreased by 38%∼71% (*P* < 0.001) after OSP treatment (Fig. 1D), but tobacco root activities improved by 46%∼172% (*P* < 0.01) after OSP treatment. Remarkably, the root activities in the 0.2% OS and 0.4% OS treatments still remained high at 15 days post-transplantation compared to those in the CK and 0.1% OS treatments (Fig. 1E). In addition, the total chlorophyll content of tobacco leaves in the 0.2% OS and 0.4% OS groups was significantly (29%, *P* < 0.01) greater than that in the CK group (Fig. 1F).

**Fig. 1.**
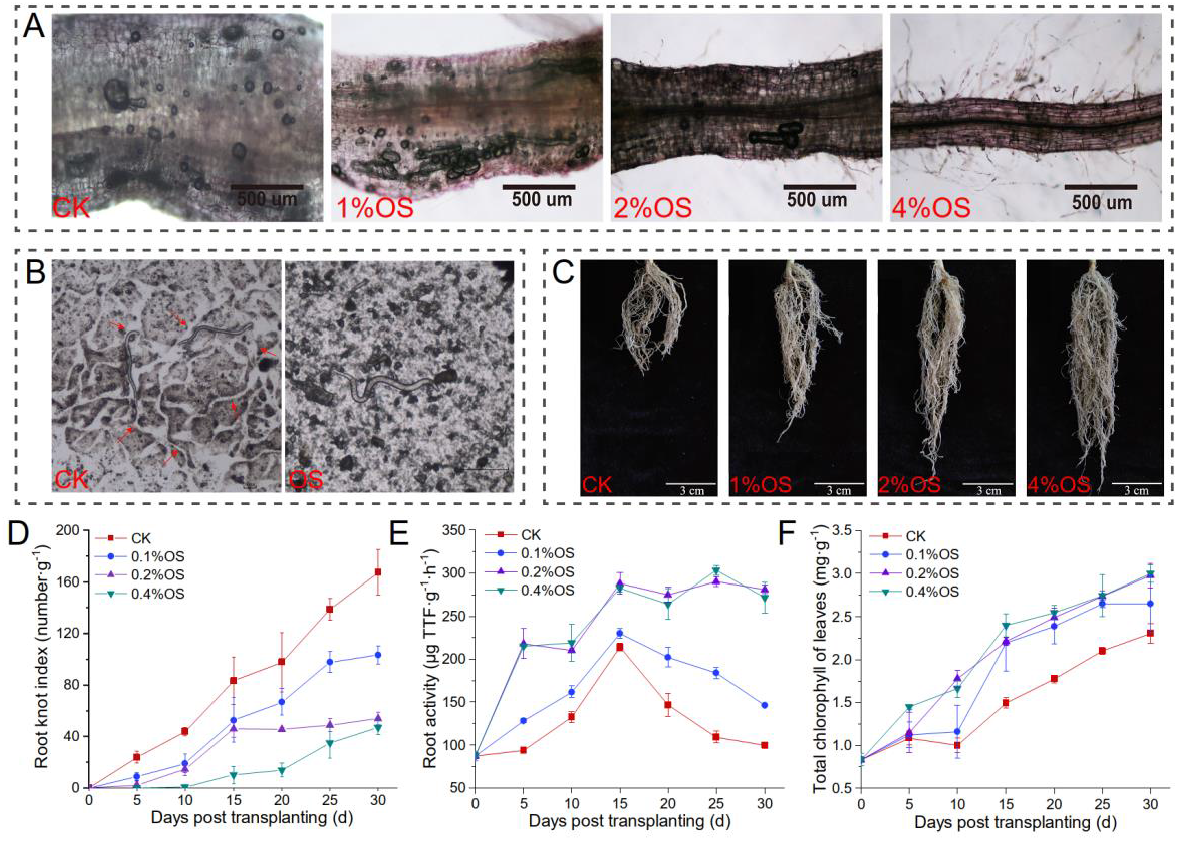
Effects of oyster shell powder on root-knot nematode infection in tobacco. (A) Root-knot nematode infection of tobacco roots 10 days post-transplantation. (B) Effects of 0.4% oyster shell powder (OS) on the migration of root-knot nematodes in soil. (C) Tobacco root growth 30 days post-transplantation. (D) Root-knot indices of tobacco roots. (E) Tobacco root activities. (F) Total chlorophyll content of tobacco leaves.

### Effects of oyster shell powder on RKN disease and soil properties in field experiments

Disease indices of RKN disease under different treatments were investigated in field experiments. The OS, AS and HB treatments significantly alleviated tobacco RKN disease in the RKN-induced field, whose disease indices were lower than those of the CK within 100 d after tobacco transplantation, and the areas under the disease process curve (AUDPC) were 56.13% (*P* < 0.01), 59.30% (*P* < 0.001) and 27.58% (*P* < 0.01) lower than those of the CK (Fig. 2A), respectively. Additionally, the OS, AS and HB treatments effectively promoted the growth of tobacco in the field (Table S4). OSP had a delayed but durable suppression effect on J2 in the soils (Fig. 2B); the density of J2 at 20 d, 80 d and 100 d post-transplantation was 15.90% (*P* < 0.01), 43.00% (*P* < 0.001) and 44.23% (*P* < 0.001) lower than that of CK, respectively. Nevertheless, the suppression effect of AS on J2 was rapid and durable, and the density of J2 was 49% (*P* < 0.001) lower than that of the CK. In addition, the suppressive effect of HB on J2 was limited.

**Fig. 2.**
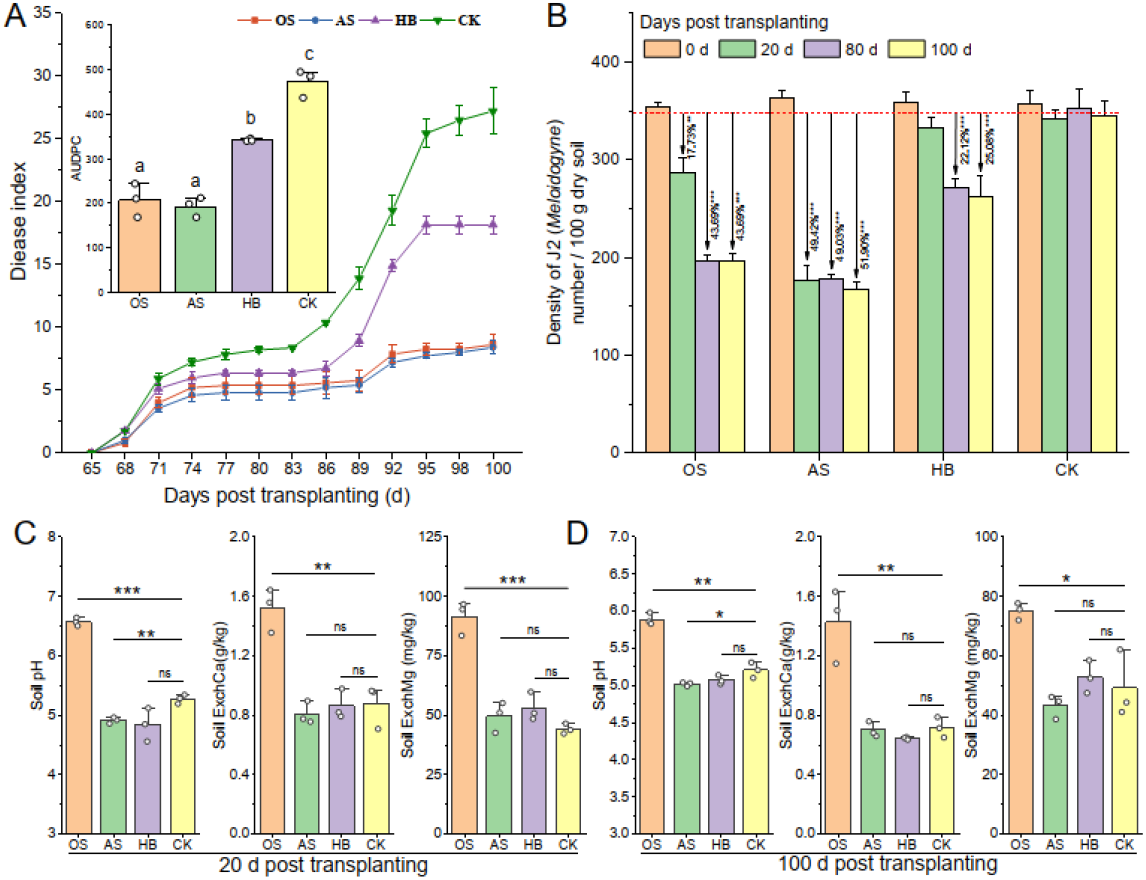
Effects of soil acidification amendment on root–knot nematode disease and soil properties. (A) Disease index curve of tobacco root-knot nematode disease and the AUDPC. (B) Density of *M. incognita* J2 in the tobacco rhizosphere. (C) Soil properties at 20 days post-transplantation. (D) Soil properties at 100 days post-transplantation.

The OSP treatment did not affect the contents of soil AvailN, AvailP, AvailK or OM within 100 d of treatment (Table S5, *P* > 0.05). Rapid and durable improvements in the soil pH, ExchCa content and ExchMg content were observed after the soil was amended with OSP (Fig. 2CD). The pH of OS increased by 0.68 (*P* < 0.01), and the contents of ExchCa and ExchMg in the OS treatment increased by 99.13% (*P* < 0.01) and 52.82% (*P* < 0.05), respectively, compared with those of the CK at 100 d post-transplantation.

### Soil microbial community composition and driving effects

Differences in the bacterial and fungal community compositions at the ASV level were detected. A total of 20715 bacterial ASVs were detected, and only 2435 of which were shared by all the treatments (Fig. 3A). A total of fungal 3472 ASVs, and only 386 of which were shared by all the treatments (Fig. 3B). The relative abundance (RA) of total fungi increased after OSP treatment, and the RA of total fungi in the OS, AS, HB and CK groups in the bacterial-fungal community were 53.35%, 50.08%, 48.71% and 51.90%, respectively (Fig. 3C). The most abundant phylum was Ascomycota, whose RA was greater than 34%, and the RA of Ascomycota in the OS group was greater than that in the other groups (Fig. 3D).

**Fig. 3.**
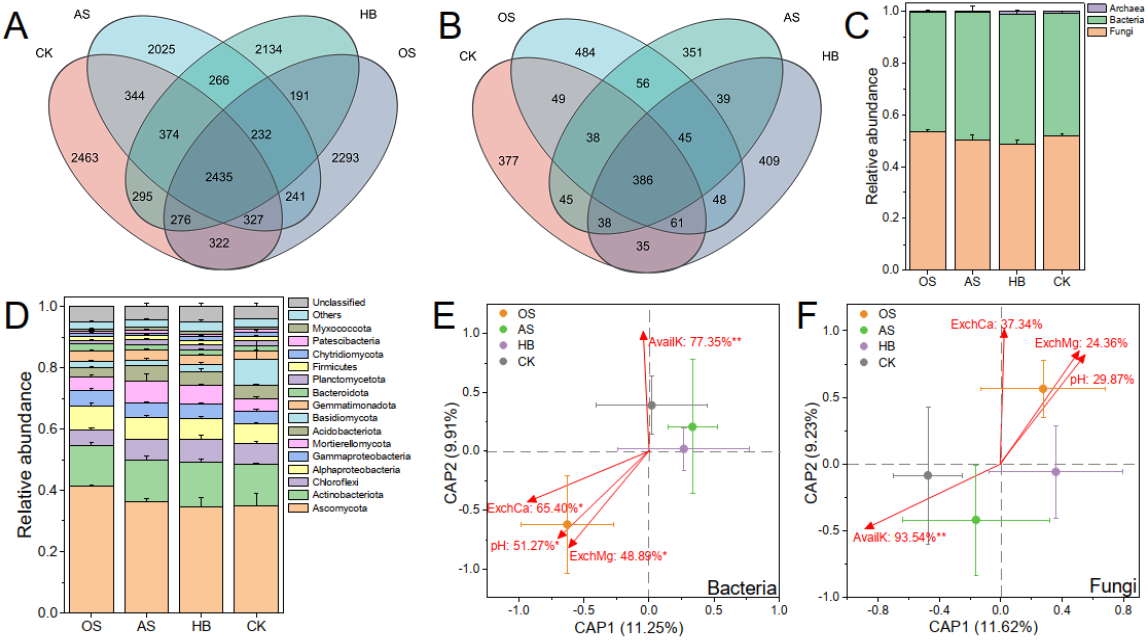
Soil microbial community composition and its driving effectors. (A) Venn diagram of bacterial ASVs of different treatments. (B) Venn diagram of fungal ASVs of different treatments. ()C Community composition at the kingdom level. (D) Bacterial–fungal community composition at the phylum level. (E) Distance-based redundancy analysis (db-RDA) between soil bacterial communities and soil properties. (F) db-RDA between the soil bacterial communities and the soil properties. Every dot represents the average of a treatment. Bars on the dots represent the standard error (SE) of a treatment. The soil properties are marked with red arrows. The text on the arrows (soil properties) represents the proportion of variance in the bacterial or fungal communities. * indicates a significant correlation, *: *P* < 0.05, **: *P* < 0.01.

The OS and HB treatments increased the richness, diversity and evenness of the soil fungal community and decreased the richness and diversity of the soil bacterial community (Table 1). The Chao1 index and Shannon index of the fungal communities in the OS treatment were greater than those in the CK by 13.65% (*P* < 0.05) and 7.77% (*P* > 0.05), respectively; moreover, the Chao1 index and Shannon index of the bacterial communities in the treatment groups were lower than those in the CK (*P* > 0.05). To reveal the key soil properties that respond to the variance in microbial communities, distance-based redundancy analysis (db-RDA) was employed. The results revealed that 36.86% of the variance in bacterial communities could be explained (*P* < 0.05) by soil AvailK (77.35%, *P* < 0.01), ExchCa (65.40%, *P* < 0.05), pH (51.27%, *P* < 0.05) and ExchMg (48.89%, *P* < 0.05) (Fig. 3E). A total of 36.01% of the variance in fungal communities could be explained (*P* < 0.05) by soil AvailK (93.54%, *P* < 0.01), ExchCa (37.34%, *P* > 0.05), pH (29.87%, *P* > 0.05) and ExchMg (24.36%, *P* > 0.05) (Fig. 3F). Our results indicate that soil ExchCa, pH and ExchMg were the most likely positive soil properties driving the assembly of bacterial and fungal communities in the OS treatment.

**Table 1.**
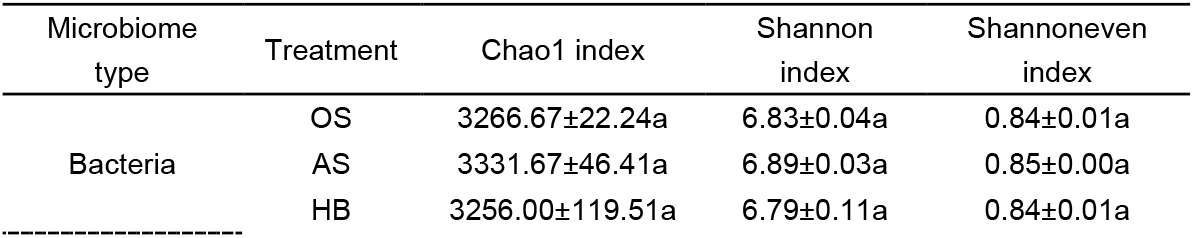
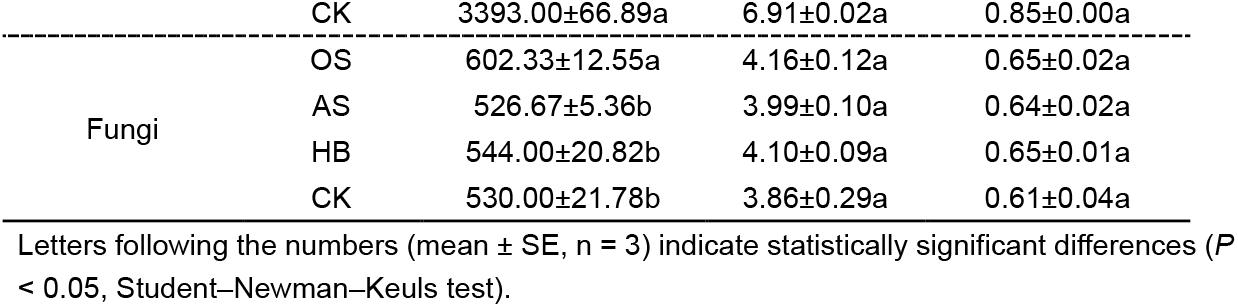
diversity of the bacterial and fungal communities of the soil samples. Letters following the numbers (mean ± SE, n = 3) indicate statistically significant differences (*P* < 0.05, Student–Newman–Keuls test).

### Key microbiomes in the fungal-bacterial co-occurrence network regulated by soil properties

A co-occurrence network with 796 bacterial ASVs and 90 fungal ASVs was constructed to reveal the interaction patterns between the bacterial and fungal communities. The results revealed that the co-occurrence network of the bacteria-fungal community was highly modular (modularity = 0.83), and eight modules (M1-M8) were identified (number of nodes ≥ 20, Fig. 4A). Bacteria tended to interact more with bacteria (79.27%), whereas fungi tended to interact more with bacteria (Fig. 4B); specifically, 68.89% of the fungi in the co-occurrence network only interacted with bacteria. The composition of interaction types (edges) in the co-occurrence network suggested that taxa tended to cooccur (positive correlations, pink lines) rather than coexclude (negative correlations, green lines), where 65.12% of the edges were positive correlations. A total of 56.04% of the interactions between bacteria and fungi were positive (Fig. 4C). Furthermore, 16 module hubs (*Z*_*i*_ ≥ 2.50, *P*_*i*_ ≤ 0.62) were identified as keystone taxa because of their important roles in network topology (Fig. 4E), 13 of which were classified into four modules, including M1 (four hubs), M2 (three hubs), M3 (three hubs), M4 (one hub) and M8 (two hubs).

**Fig. 4.**
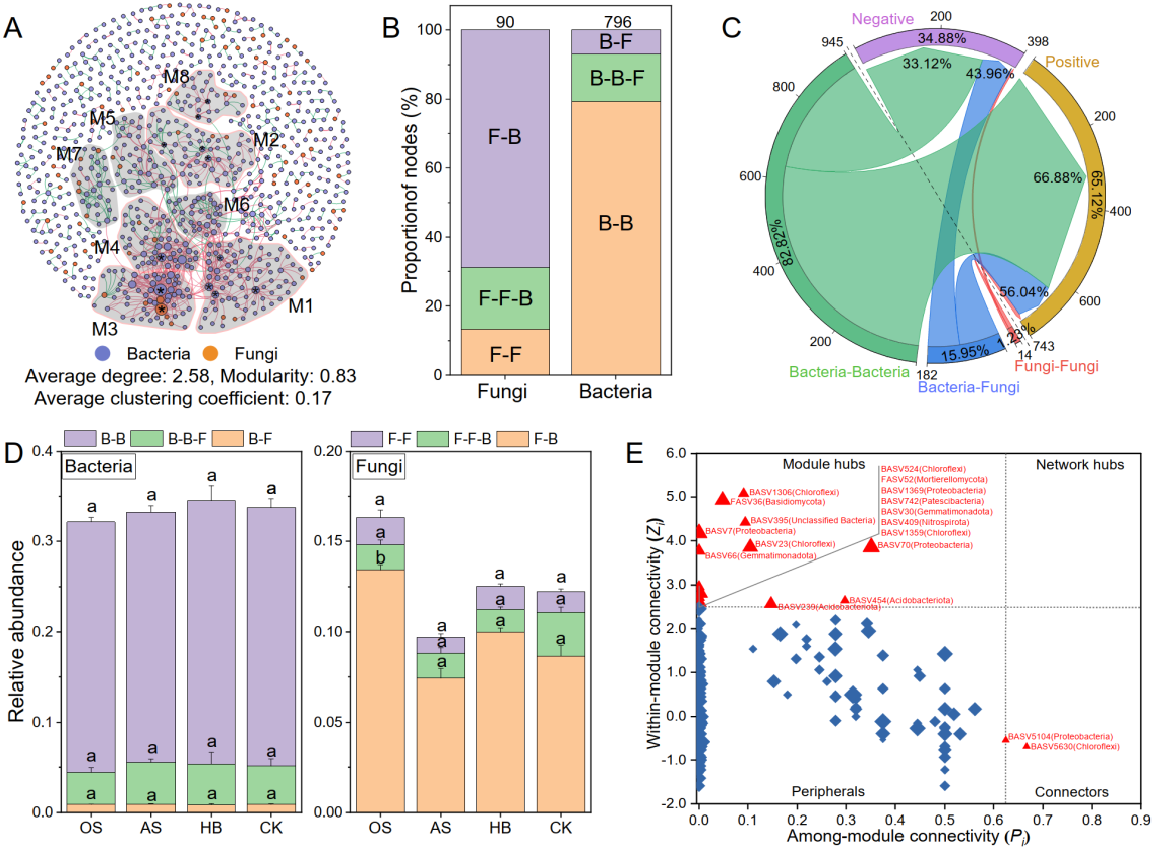
Interactions among the fungal and bacterial communities. (A) The co-occurrence network of bacterial-fungal communities at the ASV level. Nodes in the co-occurrence network are bacterial or fungal ASVs. Modules are delineated with gray areas and labeled with M1-8, representing modules with 20 or more nodes. The edge between two nodes represents a significant correlation (r ≥ 0.6, *p* < 0.05). Positive correlations are colored in pink, whereas negative correlations are colored in green. Nodes marked with * represent the identified module hubs (*Z*_*i*_ ≥ 2.50, *P*_*i*_ ≤ 0.62). (B) The composition of node types with different connections with other nodes. F-F: fungi connected with only fungi, F-F-B: fungi connected with fungi and bacteria, F-B: fungi connected with only bacteria, B-B: bacteria connected with only bacteria, B-B-F: bacteria connected with bacteria and fungi, B-F: bacteria connected with only fungi. The numbers over the charts represent the total number of nodes. (C) Composition of edge types with different correlations. (D) RA of bacterial (left) and fungal (right) nodes with different connection types in different treatments. (E) Classification of nodes to identify putative keystone ASVs within the co-occurrence networks. Each symbol represents an ASV from the network selected for detailed module analysis. The size of each symbol is related to the RA of the ASV.

M1, M2, M3, M4 and M8 of the co-occurrence network were depicted to explore the key ASVs responsive to J2 separately. The results revealed that over 80.00% of the ASVs in M2 and M8 were negatively related to J2 density and that 67.05% of the ASVs in M2 and M8 were positively regulated by OS. However, 98.63%, 98.31% and 81.13% of the ASVs in M1, M3 and M4, respectively, were positively related to J2 density, and 78.38% of all the ASVs in M1, M3 and M4 were negatively regulated by OS (Fig. 5A). Considering the close relationship between the five representative modules and J2, M2 and M8 were defined as Microcommunity S, which could suppress J2 in the tobacco rhizosphere. Moreover, M1, M3 and M4 were defined as Microcommunity C, which could benefit J2 in the tobacco rhizosphere. Additionally, Gammaproteobacteria (28.09%), Gemmatimonadota (18.67%), and Alphaproteobacteria (16.74%) were the most abundant taxa in Microcommunity S; however, Actinobacteriota (27.11%), Chloroflexi (22.42%) and Acidobacteriota (14.42%) were the most abundant taxa in Microcommunity C (Fig. 5B).

**Fig. 5.**
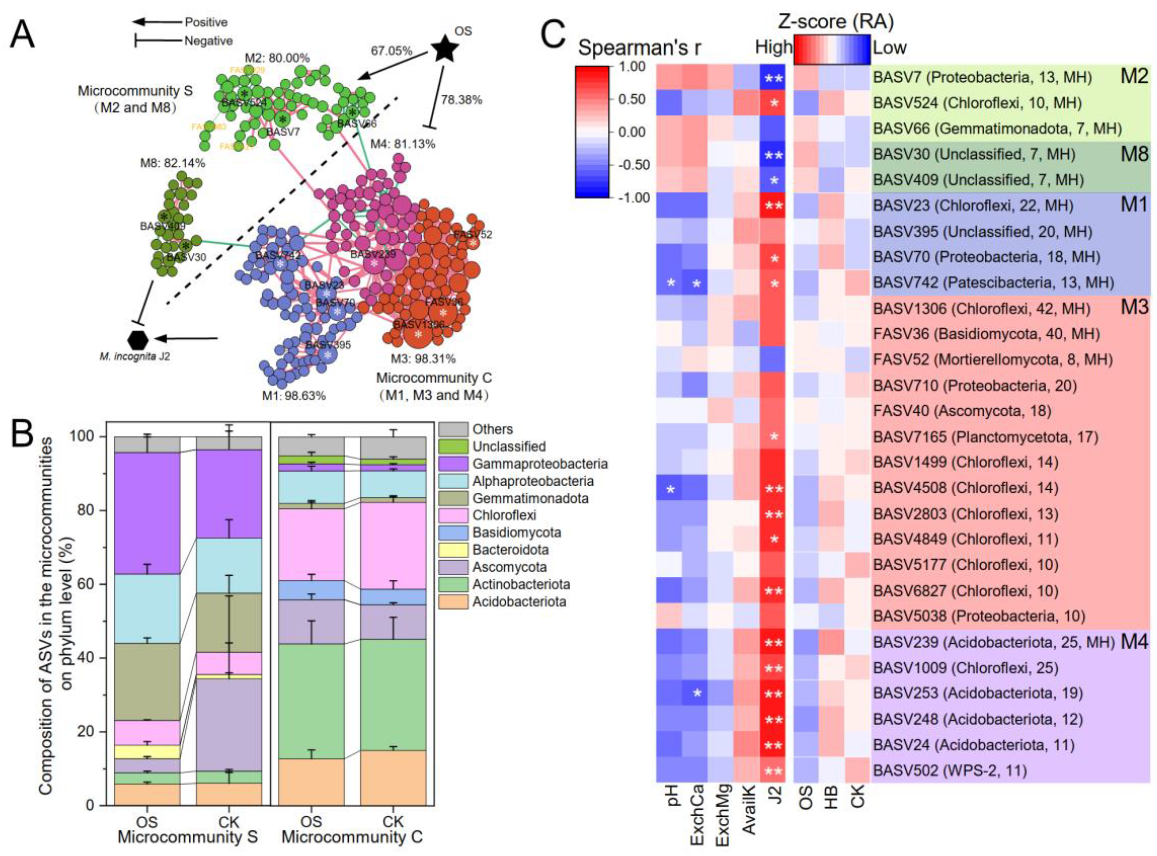
Key microbiomes related to RKNs. (A) Modules with module hubs (M1, M2, M3, M4 and M8) in the co-occurrence network are depicted. Each node represents an ASV. M2 and M8 were identified as Microcommunity S, whereas M1, M3 and M4 were identified as Microcommunity C. The numbers following the modules are the proportions of nodes in the module that are negatively or positively related to J2. Stars on the nodes mark key ASVs in the module. ASVs marked with white stars are positively related to J2 and are negatively regulated by OS. ASVs marked with black stars are negatively related to J2 and are positively regulated by OS. The numbers connecting the OS and modules are the proportions of nodes positively or negatively regulated by the OS. (B) Composition of the ASVs in microcommunity S and microcommunity C at the phylum level. (C) RA of key ASVs in the modules among different treatments and their correlations with J2 and soil properties. The labels following the ASV numbers were the classified phylum and degree of ASV in the whole co-occurrence network.

The RAs of key ASVs (module hubs or dot degrees ≥ 10) and their correlations with J2 and soil properties in Microcommunity S and Microcommunity C were analyzed (Fig. 5C). Module hubs in Microcommunity S, including BASV7, BASV66, BASV30 and BASV409, were negatively related to J2, and their RA in the OS treatment was 30.20%, 39.46%, 29.13% and 9.29% higher than that in the CK, respectively. However, module hubs in Microcommunity C, including BASV742, BASV239, BASV23 and BASV70, were positively related to J2, and their RA in the OS treatment was 75.52%, 43.00%, 19.97% and 3.99% lower than that in the CK, respectively. Additionally, 32.67% of the variance in Microcommunity S and Microcommunity C could be explained (Fig. S1, *P* < 0.05).

### Effects of soil properties and key microbiomes on Meloidogyne incognita

To quantify the effects of soil properties and the microbial community on J2, a structural equation model (SEM) was constructed with the presumed relationships among the selected subsets, including soil properties (soil pH, AvailK, ExchCa and ExchMg), the RA of Microcommunities S and C, the Chao1 index of the fungal community (Fungi.Richness) and J2 density (J2), considering that these selected subsets were least correlated while accounting for multiple drivers simultaneously. The positive and direct effects were maintained. When the soil properties pH, AvailK, ExchCa and ExchMg were considered simultaneously (Fig. S2), the SEM verified poor goodness-of-fit (χ2 = 271.884, df = 9, p = 0.000, RMSEA = 1.000, p = 0.000, CFI = 0.270), suggesting that the soil properties and microbial community characteristics were closely related to the J2 density, but the SEM needed to be modified. Hence, a new structural equation model (SEM) was constructed with the presumed relationships among the selected subsets, including soil properties (soil pH, AvailK, ExchCa and ExchMg), the RA of Microcommunities S and C, the Chao1 index of the fungal community (Fungi.Richness) and J2 density (J2), considering that these selected subsets were least correlated while accounting for multiple drivers simultaneously. The positive and direct effects were maintained. A SEM was subsequently constructed that only considered one type of soil property (Fig. 6A, Fig. S3), and the results revealed that an SEM that considered only ExchMg as the driver of the microbial community was verified to have a strong goodness-of-fit (χ2 = 7.713, df = 9, *p* = 0.563, RMSEA = 0.000, *p* = 0.579, CFI = 1.000). Fungi.Richness and Microcommunity C explained 93% of the variation in J2 density (Fig. 6A); specifically, J2 density was negatively affected by Fungi.Richness (Standardized total effect (STE) = -0.64, *p* < 0.001) and was positively affected by Microcommunity C (STE = 0.50, *p* < 0.001). Furthermore, soil ExchMg, which was positively regulated by the OS treatment (STE = 0.96, *p* < 0.001), had positive effects on Microcommunity S (STE = 0.57, *p* < 0.05) and Fungi.Richness (STE = 0.82, *p* < 0.001). Additionally, the total effects (direct and indirect) of OS, ExchMg, Microcommunity S, Microcommunity C and Fungi.Richness of J2 were -0.75, -0.78, 0.64, -0.44 and 0.50, respectively (Fig. 6B). Overall, OSP promoted microbial-mediated suppression of J2 by increasing the content of soil ExchMg in the tobacco rhizosphere.

**Fig. 6.**
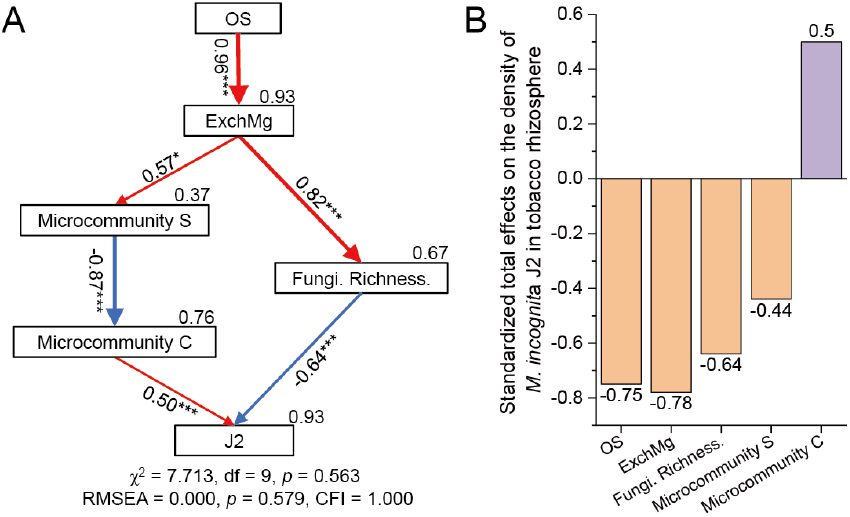
Biotic and abiotic drivers regulating the density of *M. incognita* J2. (A) Structural equation model (SEM) showing the influences of the OSP treatment, soil properties and microbial profiles on J2. The numbers near the pathway represent the standardized path coefficients. Bootstrap-based *p* values for path coefficients are indicated by *** when *p* < 0.001, ** when *p* < 0.01 and * when *p* < 0.05. The numbers on the rectangles represent the proportion of variance explained. χ^2^: Chi-square; df: degrees of freedom; = 9, *p*: probability level; RMSEA: root-mean-square error of approximation; CFI: comparative fit index. (B) Standardized total effects (direct and indirect effects) based on SEM.

## DISCUSSION

Considerable progress has been made in the use and development of bioresources as a strategy to manage PPNs. Cole *et al*. reported that the application of biochar could decrease the populations of plant-parasitic nematodes and increase the abundance of predatory nematodes (20). In this study, we reused oyster shell as a natural bioresource and discovered the ability of oyster shell powder to suppress second-stage juveniles of *Meloidogyne incognita* (J2) for the first time. Calcinated oyster shell powder (OSP) reduced the tobacco root-knot index and J2 density by inhibiting the migration of J2s and enhancing the microbially mediated suppression of J2s in the tobacco rhizosphere. Moreover, we revealed the positive involvement of soil properties, especially soil exchangeable magnesium (ExchMg), in enhancing microbial-mediated suppression of J2; specifically, Chloroflexi- and Acidobacteriota-dominated microbes promoted RKN prosperity, and Proteobacteria- and Gemmatimonadota-dominated microbes and fungal richness contributed to the suppression of RKNs in the tobacco rhizosphere. On the basis of these results, we believe that calcinated OSP has the potential to be developed as a bioresource to suppress *Meloidogyne incognita*.

China produced approximately 80% of oysters worldwide in 2015 (21), and wasted oyster shells were improperly discarded and accumulated, thereby presenting a threat to public health in recent decades (22). Therefore, the reuse of wasted oyster shells in crop production has practical value for natural bioresource recycling, eco-friendly crop protection and sustainable agriculture. Previous studies have reported that alkaline OSP contains abundant and high-dissolutive nutrients, including Ca and Mg (23), and this explanation is supported by the finding that the application of calcinated OSP majorly increased the soil pH and contents of soil ExchCa and ExchMg in our study. Notably, nutrients released by OSP have several benefits for plant growth promotion (23) and soil microbiome-mediated disease control (10). Yang et al. reported that OSP promoted the grain weight of rice by ameliorating growth inhibition caused by Cd toxicity (24). The promotion of agronomic growth in OSP-pretreated tobacco was consistently observed in our field experiments (Table S4). Zhang *et al*. revealed that the CaO-mediated recruitment of potentially beneficial rhizobacteria enhances disease suppression of bacterial wilt by alleviating soil acidification (25, 26). Cao et al. confirmed that the pH level and the levels of calcium (Ca), magnesium (Mg), phosphorus (P) and iron (Fe) were key environmental indicators influencing the composition of the microbial community in the tobacco rhizosphere during RKN infection (27). Our findings suggest for the first time the indirect and positive effects of OSP on RKN suppression in the tobacco rhizosphere, which are mediated by soil ExchMg and the microbial community.

The rhizosphere microbiome provides a first line of defense for plants against RKNs (28, 29). Cao *et al*. reported that the richness and diversity of the fungal community in the healthy group were significantly greater than those in the RKN infection group (3), and our finding via SEM that J2 density was negatively affected by fungal richness is consistent with their results (Fig. 6). Notably, some specific fungi are natural enemies of nematodes and have direct or indirect effects on nematode inhibition (30). The OSP-promoted increase in fungal richness may result in an increase in nematophagous fungi, thereby reducing the RKN population (31). Moreover, our findings indicate that Chloroflexi and Acidobacteriota might play important roles in RKN prospering in the tobacco rhizosphere; in contrast, Proteobacteria and Gemmatimonadota might contribute to RKN inhibition (Fig. 6B). Previous studies have suggested a relationship between the four kinds of bacterial groups and RKNs. Jiang *et al*. reported that nematodes enriched certain soil microbiome groups, including Chloroflexi and Actinobacteria, in wheat fields (32). These studies partly support our identification of Chloroflexi and Acidobacteria as RKN disease-induced microbiomes. There are few reports about the relationship between Gemmatimonadota and RKNs, but Li et al. reported a potential plant growth-promoting function of Gemmatimonadota, as a significant increase in strawberry yield was positively correlated with increases in Gemmatimonadota (33). Mugnai et al. revealed that Gemmatimonadota could promote both inter- and intra-kingdom interactions in the soil microbial community (34), and these characteristics of Gemmatimonadota partly support our identification of it as a disease-suppressing microbiome. However, the relationships of Proteobacteria with RKNs identified in previous studies have been inconsistent. For example, Zheng et al. reported that the diversity of the nematode *Dorylaimus stagnalis* was negatively correlated with the abundance of Proteobacteria in rice fields (35), but Castillo et al. reported that alpha-Proteobacteria was positively correlated with *Meloidogyne chitwoodi* (36). Therefore, the effects of Proteobacteria on RKN disease still need to be investigated.

Our findings highlight the microbial-mediated suppression of *Meloidogyne incognita* and the significant regulatory role of magnesium ions in microbial-mediated disease suppression. However, our results on the effects of microcommunities on RKNs and the regulatory role of magnesium ions rely on sequencing data and correlation analysis, and we believe that experimental validation, such as field trials or greenhouse experiments, to demonstrate these direct effects in the future would further confirm the findings and make our conclusions more robust. Moreover, evaluation of the potential control effect of OSP on other kinds of PPNs would make it more applicable in crop protection.

## Materials and methods

### Pot experimental design

The pot experiment was conducted from January 1^st^ to February 25^th,^ 2022, at the College of Plant Protection of Southwest University, Chongqing, China. Soil collected from a field with RKN disease (*Meloidogyne incognita*) was used in the pot experiment. The field is located in Liangshan Yi Autonomous Prefecture, Sichuan Province (26°17′38″N, 102°01′10″E, elevation: 1892 m) and has been subjected to continuous tobacco cropping for several years. The physiochemical characteristics and information of oyster shell powder (OSP) used in the pot experiment is shown in Table S1 and S2 in the Supplementary material (10). Honghuadajinyuan (*Nicotiana tabacum* L., susceptible tobacco), bred and cultured for 30 days under identical conditions, was used in the pot experiment.

The OSP and soil were mixed at different mass ratios and used in the pot experiments. There were four treatments, each treatment had four repetitions, and every repetition had 30 tobacco plants. The four treatments were as follows: (1) 0.1% OS: a mass ratio of OSP to soil of 0.1%; (2) 0.2% OS: a mass ratio of OSP to soil of 0.2%; (3) 0.4% OS: a mass ratio of OSP to soil of 0.4%; and (4) CK: a blank control, without any treatment. The mixtures or soils of different treatments were cultured for two weeks under identical conditions before tobacco transplantation.

### Field experimental design

The field experiment was conducted from May 1^st^ to August 25^th^ in 2022 in the RKN disease-induced tobacco field mentioned above. Yunyan87 (*Nicotiana tabacum* L., susceptible tobacco) was used in the field experiment. Oyster shell powder, 3% avermectin·fosthiazate (0.5% avermectin and 2.5% fosthiazate) and 0.25 billion live spores/g *Verticillium chlamydosporium* were used in the field experiment (Table S1 in the Supplementary material).

Four treatments were set in the field experiment, and the four treatments were as follows: (1) OS: Oyster shell powder, 100 g/plant mixed with soil as a nest fertilizer before tobacco transplantation; (2) AS: 3% avermectin·fosthiazate, 2 g/plant mixed with soil as a nest fertilizer before tobacco transplantation; (3) HB: 0.25 billion live spores/g *Verticillium chlamydosporium*, 2 g/plant, irrigating just after tobacco transplantation; (4) CK: a blank control without any treatment. and each treatment had three repetitions (plots). Each plot was planted with 100 plants, consisting of four 15 m-long rows spaced 1.2 m apart, with a spacing between two close plants in one line of 0.55 m. All fertilizers used in the tobacco field were applied once as the base manure before tobacco was transplanted.

### Data investigation

Tobacco roots in the pot experiment were cleared and stained with acid fuchsin 10 days post-transplantation (37) and then photographed under an inverted fluorescence microscope (Nikon, Japan). Tobacco root–knot indices (number of root-knots per gram of tobacco roots) were investigated every 10 days for 30 days (38); RKN in the 0.4% OS and CK soils were photographed under an inverted fluorescence microscope; Tobacco root activities were investigated by colorimetry every 10 days for 30 days (39), and the total chlorophyll content of tobacco leaves was investigated by colorimetry every 10 days within 30 days in the pot experiment (40).

RKN disease grade of tobacco plants aboveground was investigated every 3 days after the first symptomatic day in the field experiment. The criteria of disease grade referred to the national standard of the tobacco pest classification survey method of China (Supplementary material Table S3) (41). Disease index and area under disease progress (AUDPC) were calculated (42,43).

### Soil samples and sample tests

Soil samples from the bulk soil and tobacco rhizosphere were collected at a sampling depth of 10–20 cm at 0, 20, 80, and 100 days after transplanting in the field experiment, which represented different periods of disease progression. The collection method for the soil samples followed those of our previous study (44). Five samples of representative bulk soil and three rhizosphere soil samples for each treatment were collected in the experimental field (three repetitions/plot).

The density of second-stage juveniles of *Meloidogyne incognita* (*M. incognita* J2/J2) in each soil sample at 0, 20, 80 and 100 days after transplantation was determined. Hatched J2 in the soil samples were isolated via the Baermann funnel method, and the populations were counted using a stage micrometer under a Nikon microscope (Tokyo, Japan). The density of J2 was converted to a number per 100 g dry soil (6).

The properties of the soil samples were measured at 20 and 100 days after transplanting. Soil pH, organic matter (OM), available nitrogen (AvailN), available phosphorus (AvailP), available potassium (AvailK), exchangeable calcium (ExchCa) and exchangeable magnesium (ExchMg) were measured using standard methods. In particular, soil pH was measured using a pH electrode (Metter-Toledo SevenMulti™, Switzerland) at a soil: water ratio of 1:2.5 (w/v). OM was determined using the Kjeldahl digestion method (45) and the dichromate oxidation method (46). AvailP was extracted with 0.025 mol l^−1^ HCl + 0.03 mol l^−1^ NH4F and measured by a visible spectrophotometer (47). AvailK, ExchCa and ExchMg were determined in ammonium acetate extracts by flame photometry (48).

### Sequencing

Microbial DNA from the soil samples was extracted using the FastDNA Spin Kit (MP Biomedicals, United States kits) following the standard protocol. The amplification and purification of soil microbial DNA were conducted according to the methods described in our previous study. ITS1 forward and ITS2 reverse primers were used to amplify the ITS1 region of the fungal ITS gene. The 515 forward and 806 reverse primers were used to amplify the V4 region of the bacterial 16S rDNA gene (8).

The sequencing of 16S rRNA and ITS gene fragments was conducted by Shanghai Majorbio Co., Ltd., China. using the Illumina MiSeqPE250 platform. The quality control and annotation of the raw sequencing data adhered to the methods described in our previous study (8). Amplicon sequence variants (ASVs) were assigned to taxa at the phylum, class, order, family and genus levels via QIIME2. The Silva138 database was used for the taxonomic assignment of bacterial and archaeal ASVs, and the Unite 8.0 database was used for the taxonomic assignment of fungal reads.

### Data analyses

Microbial apha diversity and beta diversity analyses were performed via the Majorbio Cloud Platform (www.majorbio.com). Specifically, α diversity indices of bacterial and fungal communities were calculated on the basis of faith phylogenetic metrics at the ASV level. Principal coordinate analysis (PCoA) and distance-based redundancy analysis (db-RDA) was performed on the basis of the Bray-Curtis distance according to the phylogenetic tree to determine the dissimilarity of the microbial communities. The underlying co-occurrence network among bacteria and fungi was depicted at the ASV level (relative abundance over 0.01%) through network analysis via the Molecular Ecological Network analysis pipeline (49). ASVs represented in more than 50% of the samples were reserved, and data filtering was performed prior to avoiding zero values that could result in spurious correlations (50). A structural equation model (SEM) was constructed in IBM SPSS AMOS software. The statistically significant results were assessed by ANOVA in SPSS (Statistical Product Service Solutions). Linear regression analysis was executed in SPSS to evaluate the correlation between variables. Other figures were plotted via Origin software.

## ACKNOWLEDGMENTS

This work was supported by the Key Project from Sichuan Company of China National Tobacco Corporation (202051340024416 and SCYC202010).

Q.P.J., Y.W., W.D. and S.L.L. conceived and designed the study. Q.P.J., J.M.Y. and D.J.X., performed the analyses. J.F.W., L.Q.J., D.Y.L. and Q.D. performed the investigation and sampling. Q.P.J. drafted the manuscript. Y.Q., S.P.G. and X.W.Y. made plentiful valuable comments for the manuscript. All the authors reviewed and approved the manuscript.

## FUNDING

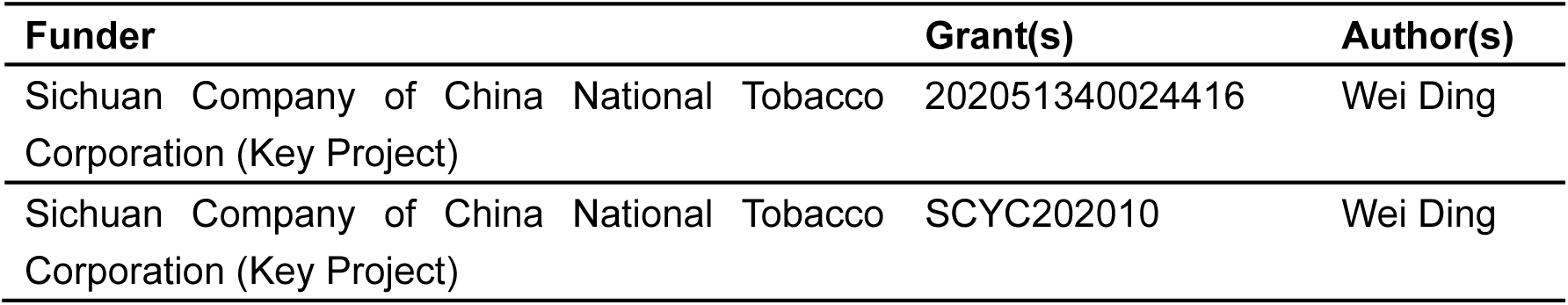

## DATA AVAILABILITY

The data that support the findings of this study are openly available in the NCBI Sequence Read Archive (SRA) database under accession number PRJNA1080250 for 16S rRNA, and PRJNA1080272 for ITS.

## SUPPLEMENTAL MATERIAL

Supplemental material may be found in.

